# Stage-specific screening reveals complex resilience pathways to cold stress in rice

**DOI:** 10.1101/2025.11.23.690040

**Authors:** Fahamida Akter, Partha S. Biswas, A.K.M. Aminul Islam, M.S. Raihan, Md. Mizanur Rahman, K. M. Iftekharuddaula, Mohammad Rafiqul Islam, John Damien Platten

## Abstract

Rice cultivation in Bangladesh’s northern and northeastern regions face a critical challenge – cold stress during the seedling and reproductive stages, which can drastically reduce yield. As a precursor to generate improved elite lines, a diverse panel of rice germplasm was screened to identify genotypes with resilience to low temperature at both key developmental phases. Seedling-stage tolerance was assessed using an artificial cold-water tank, while reproductive-stage tolerance was evaluated under both natural field conditions and controlled cold screening facilities. Performance was measured through a combination of quantitative traits (plant height, days to heading, panicle length) and qualitative indicators (panicle degeneration, panicle exsertion, spikelet fertility). Two breeding lines - BR8907-B-1-2-CS1-4-CS2-P3-4 and BR8909-B-12-2-CS1-4-CS2-P2-3-2, demonstrated seedling-stage cold tolerance with minimum leaf discoloration and the highest survival rate across three experimental batches. Five genotypes (Bhutan, IR83222-F11-173, Rata Boro, BRRI dhan74, BR11712-4R-227) showed tolerance during the reproductive stage. Three lines (Bhutan, BR11712-4R-227, and BR12266-44-11-32-5-1-1-HR10-B) showed moderately tolerant to tolerant across both stages. Six genotypes, including BR10317-5R-25, IR18A1859, and BRRI dhan28 were consistently vulnerable to cold stress at both stages. Principal Component Analysis (PCA) revealed that under field conditions, both seedling and reproductive traits contributed to shared components - suggesting overlapping physiological mechanisms. However, under controlled environments, the traits separated distinctly, pointing to stage-specific genetic control. These results are consistent with reports of distinct QTLs contributing to cold tolerance at different stages, highlighting the complex nature of cold tolerance and informing breeding strategies for enhanced cold resilience in rice.

## Introduction

For more than half of the world population, rice is used as a staple food, cultivated in more than 100 countries. With the global population projected to reach 11 billion by 2100 [1], having already exceeded 7.5 billion people, ensuring sufficient rice production is paramount. In Bangladesh, rice accounts for 97% of total food grain production for its 172.92 million people [2]. By 2050, Bangladesh’s population is expected to hit 215.4 million, requiring an estimated 44.6 million tons of milled rice [3]. However, decreasing arable land and increasing climate vulnerabilities such as drought, salinity, floods, heat, and waterlogging threaten food security. Among these, low-temperature stress (LTS) significantly limits plant growth, productivity, and yield affecting 10% of the 130 million hectares of rice globally [4]. Rice grown in tropical and subtropical Asia during winter is particularly sensitive to cold stress [5], especially during the seedling and reproductive stages [6,7]. In Bangladesh, approximately 2 million hectares of rice are impacted by LTS [8].

During the winter season in northern Bangladesh, temperatures drop below 10°C at the seedling stage, causing leaf discoloration, yellowing, rolling, and stunted growth, often leading to seedling mortality and increased production costs [9]. Cold stress at the reproductive stage causes abnormality at anthesis, resulting in the cessation of anther development, immature pollen grains, non-emergence of anthers from florets, anther indehiscence, pollen grains remaining in anther loculi, poor pollen shedding, and failure of pollen to germinate after reaching the stigmas. These cause panicle degeneration, incomplete panicle exsertion, and spikelet sterility, which ultimately lower the grain yield of rice [7,10–12]. Reproductive-stage cold stress thresholds are 20°C and 15°C for cold-sensitive and cold-tolerant varieties, respectively [13]. Flash floods cause catastrophic damage to Boro season rice in the northeastern Haor region of Bangladesh. To escape flash floods, early sowing of Boro rice is recommended [14]. However, this causes the booting stage to coincide with low-temperature stress (15-19°C; [8]) with spikelet sterility sometimes reaching 100%. Therefore, developing short-duration, high-yielding, cold-tolerant rice varieties is crucial to overcome the dual threats of flash floods and cold injury. To develop a variety, availability of cold-tolerant donor parents is a prerequisite. Thus, the availability of improved methods for germplasm assessment is of high value for breeding programs [15]

Screening of rice genotypes for cold tolerance under only natural field conditions may not be efficient because the natural climate is unpredictable in terms of its intensity, duration, and timing. The success rate of screening cold tolerance in natural field conditions is very low. This may be due to the complexity of cold stress tolerance traits, low genetic variance of yield components due to stress, and lack of proper selection criteria [16]. On the other hand, screening conducted under controlled (artificial) conditions may perform differently from actual field conditions. It’s therefore desirable to consider both field and artificial conditions when screening for tolerant genotypes [17]. This approach facilitates the large-scale identification and selection of cold-tolerant genotypes. Previous researchers used a reliable method of phenotyping for cold tolerance under both natural and artificial conditions [7]. The objective of this study was to screen for cold-tolerant genotypes by evaluating their performance at two key stages: the seedling stage, under controlled (artificial) conditions, and the reproductive stage, under both natural and artificial conditions.

## Materials and methods

### Plant materials

A diverse panel of 46 rice genotypes, including advanced breeding lines, local landraces, and BRRI varieties, was collected from the Plant Breeding Division, BRRI, Gazipur. The materials were evaluated for cold tolerance at the seedling and reproductive stages. In cold screening at the seedling stage, 34 genotypes and four check varieties were tested, whereas 23 genotypes and three check varieties were evaluated for reproductive stage cold screening. Among them, 18 genotypes are common in both seedling and reproductive stage cold screening (**Table 1**).

**Table 1:**
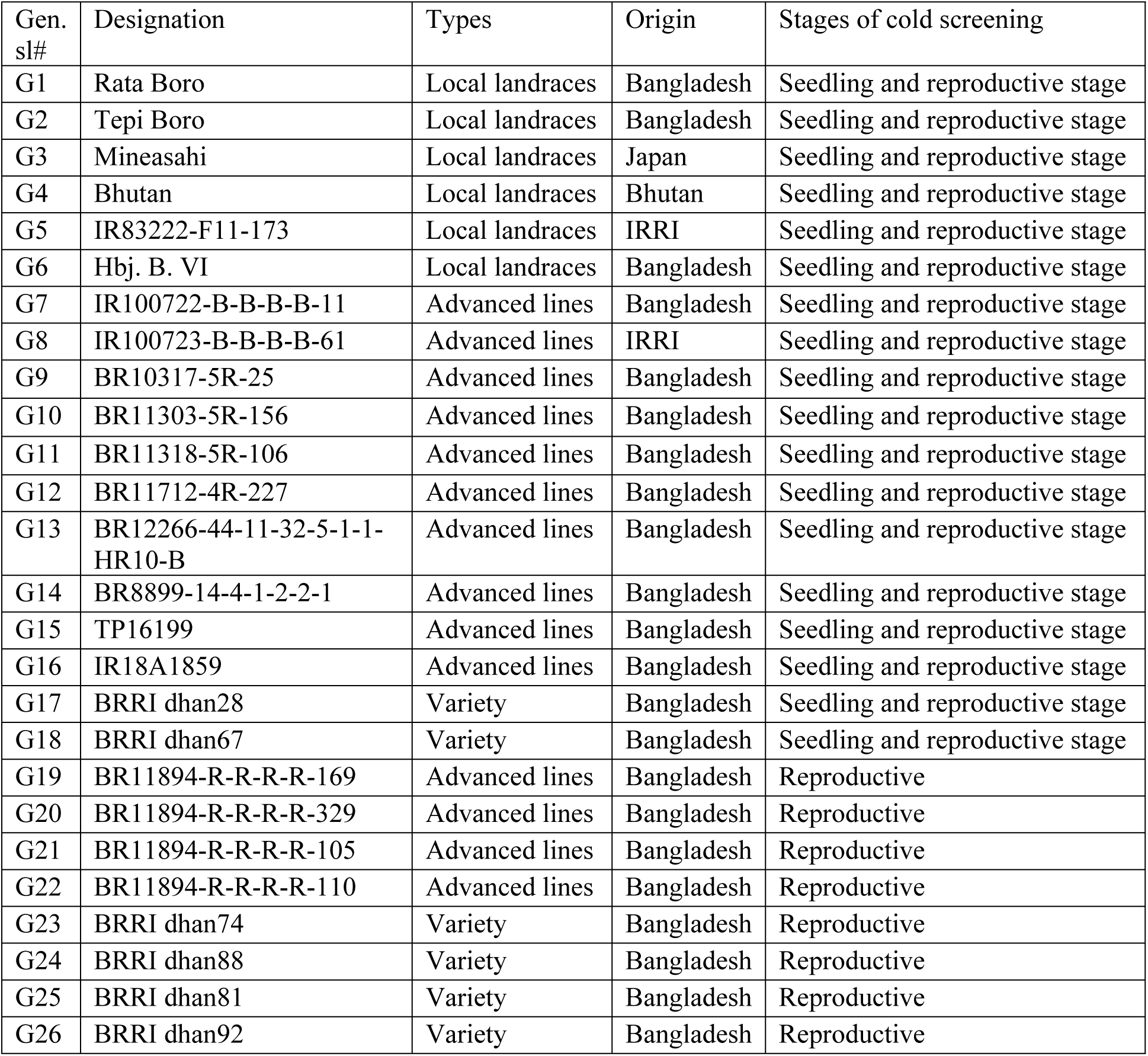
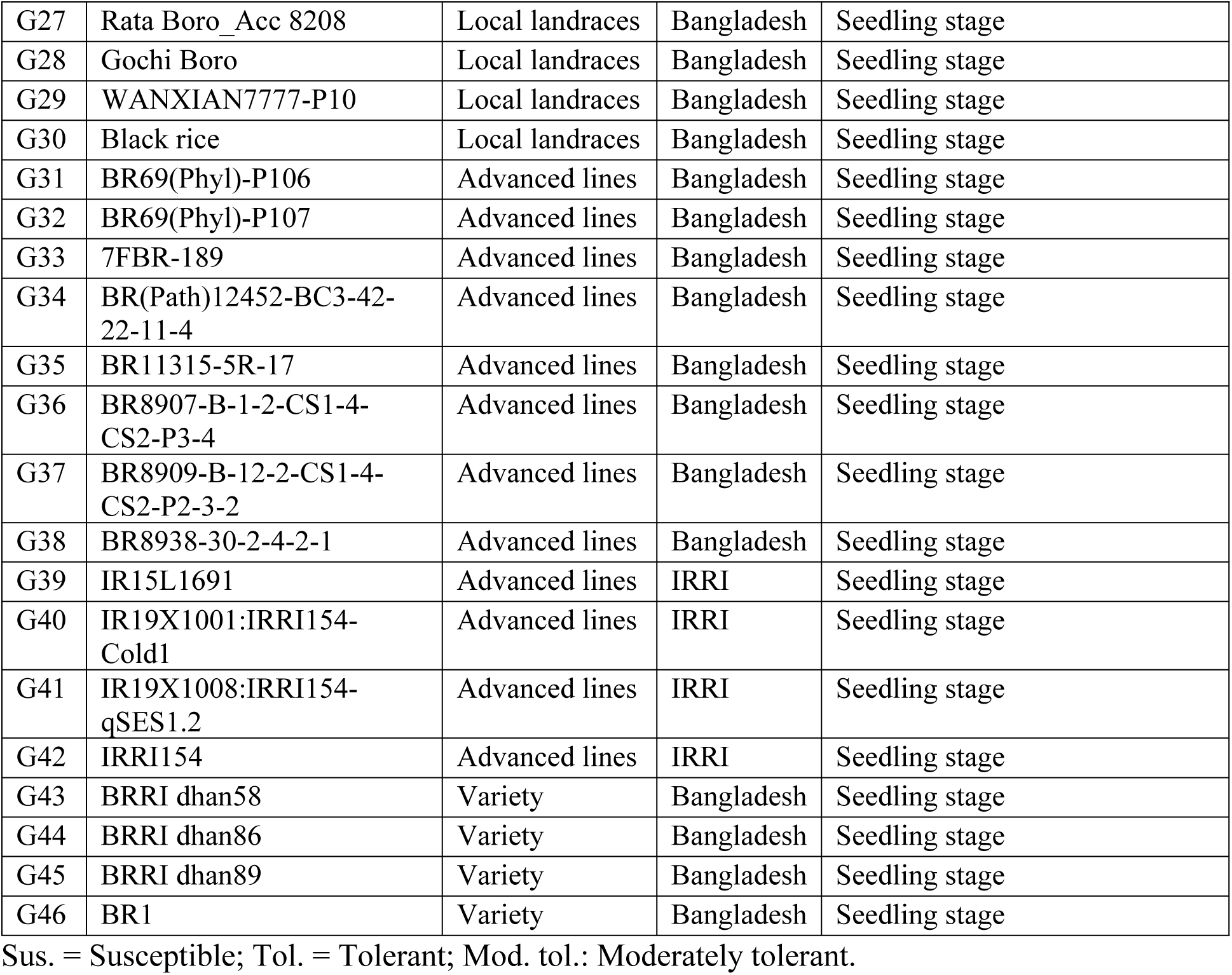
List of genotypes used for cold screening.

### Screening of cold tolerance at seedling stage

A total of 34 breeding lines were evaluated under artificial cold stress conditions at **13°C**, using cold-water irrigation in a tank setup[18]. BRRI dhan67 and HBJ B VI were used as cold-tolerant check varieties, while BR1 and BRRI dhan28 were included as susceptible check varieties [19]. Each genotype, including the checks, was represented by 10 seedlings planted in single-row plots with 3.0 cm spacing. The seedlings were grown in flat-bottomed plastic trays (dimensions: 60 cm × 30 cm × 2.5 cm) containing fertilized soil free of crop residues and gravel. A total of 34 breeding lines, along with four check varieties, were evaluated using a row column design. The experimental units were arranged in trays, each of which was subdivided into two blocks, serving as blocks within the replication.

The experiment was conducted in three batches (Batch-1, seed sown on 6/22/2022; Batch-2, seed sown on 7/30/2022; Batch-3, seed sown on 7/24/2024) with four replications in Batch-1 and Batch-2 and six replications in Batch-3. A thin layer of finely granulated soil was applied after sowing to cover the germinated seeds. The seedlings were allowed to grow until the 3-leaf stage (8-12 days), and then the trays were placed into the cold-water tank [20] that was pre-set at 13°C. In Batch-1, the temperature in the cold-water tank was not stable throughout the tank. The water temperature at six positions (4 points at the four corners and two points across the middle of the tank) within the tank ranged from 10.3 to 14℃. Then, an aquarium internal filter motor (model: SOBO WP-1200F) of AC 220V 50Hz 15 watts was installed into the tank to circulate the movement of water to stabilize the temperature close to 13℃ all over the tank. The temperature was recorded five times daily with an interval of 3 hours (6 AM, 9 AM, 12 PM, 3 PM, 6 PM) within the period of cold treatment (**S1 Fig**). The duration of cold treatment across three batches ranged from 8 to 14 days. The water level was maintained at 2 cm above the soil level to keep the trays submerged. Cold stress symptoms were assessed using a subjective leaf discoloration (LD) scale ranging from 1 to 9, following the protocol outlined by Shim et al. (2022) [21] (**Table 2**). LD scoring was conducted at the point when the susceptible check variety, BR1, exhibited complete mortality; normally this is done within six to ten days after cold treatment starts (Batch 1: 6 days, Batch 2: 14 days, Batch 3: 10 days) [19]. After recording the LD score, the trays were placed at ambient temperature in a sunny place for recovery from cold stress. The survival count was recorded from each plot after a seven-day recovery period. The survival rate in percentage was calculated as the percentage of recovered green plants to the total number of plants used in the cold treatment.

**Table 2:** Leaf discoloration score for cold tolerance at seedling stage.

| Scale for cold tolerance according to SES |  |
| --- | --- |
| 1 | No damage to leaves, normal leaf color (0-10% leaf tip dried) |
| 3 | Tip of leaves slightly dried, folded and light green, 11-30% leaf tip dried |
| 5 | Seedlings moderately folded and wilted, 31-50% seedlings dried, pale green to yellowish leaves |
| 7 | Seedlings severely rolled and dried; reddish-brown leaves (susceptible), 61-80% seedlings dried |
| 9 | Most seedlings dead and dying (highly susceptible), 81-100% seedlings dried |

### Screening of cold tolerance at the reproductive stage

Screening of 22 diverse rice genotypes was carried out for reproductive stage cold tolerance along with three standard check varieties: BRRI dhan28 as susceptible, and BRRI dhan67 and IR83222-F11-173 as a moderately tolerant check and tolerant check, respectively. Screening was conducted under two different conditions: natural field conditions during the Boro season and artificial cold screening facilities (Phytotron) under cold stress and non-stress conditions.

### Field condition (natural condition)

To carry out field screening, one set of genotypes was sown on 21 October 2024 (34 days earlier than the regular sowing). This ensured the panicle initiation (PI) to booting stage of the crop would be exposed to low temperature from the end of January to the first week of February. This was the cold stress plot. A second set of the same genotypes was sown in the regular Boro season on 25 November and was treated as a non-stress plot. Flowering window and temperature profile of 25 genotypes were presented in **S2 Fig** and **S3 Fig** respectively, for cold stress (A) and non-stress (B) conditions. Data on plant height, effective tiller number, days to heading, and panicle length were measured; vegetative stage score (VegS), panicle degeneration score (PDS), panicle exsertion score (PES), and spikelet fertility score (SFS) were recorded based on SES (Standard Evaluation System) presented in **Table 3**. The rate of reduction (RR) was calculated by the following method: RR = {(Mean value of nonstress - Mean value for cold stress)/mean value of non-stress} × 100% [17].

**Table 3:** Scoring details on cold injury-related traits.

| Score | Data collected | Scoring scale |
| --- | --- | --- |
| Vegetative stage | Maximum tillering stage | 1 = Excellent, 3 = Good, 5 = Fair, 7 = Poor, 9 = Very poor: based on growth, tiller number and leaf color |
| Panicle degeneration | Flowering stage | 1 = 0%; 3 = 1-10%; 5 = 11-25%; 7 = 26-40% 9 = >40% of panicle degenerate |
| Panicle exertion | Hard stage Dough stage | 1 = >1.0 cm, 3 = 0.5 - 1.0 cm, 5 = 0.0 - 0.5 cm, 7 = <0.0 cm - 1/4th of the panicle, 9 = >1/4th of the panicle length |
| Spikelet fertility | Hard stage Dough stage | 1 = Highly fertile (>90%), 3 = Fertile (75-89%), 5 = Partly sterile (50-74%), 7 = Highly sterile (50% to trace), 9 = 0% |

### Cold screening in Phytotron

The same set of genotypes were evaluated in a artificial cold screening facilities (phytotron) with controlled air (15.9-19.14℃, average 17℃) and water (16.66-20.24℃, average 18℃) conditions, considered as cold stress (**S4A Fig**) following a RCB design with 3 replications. The phytotron setup consisted of two water tanks, wherein a total of 100 entries can be accommodated for cold screening at the reproductive stage. Thirty-day-old seedlings of each genotype were transplanted into six plastic pots with a single seedling per pot. Three pots were treated as cold stress, and three pots were used as a non-stress control. Plants were grown in ambient temperature in the net house until the meiotic phase of the reproductive stage began. The reproductive stage was determined by the distance between the ligule of the flag leaf and that of the penultimate leaf (Yoshida, 1981), considering an interval of -3 (flag leaf ligule below the penultimate leaf ligule) to +10 cm (flag leaf ligule above the penultimate leaf ligule) as indicative of this stage [19]. In the reproductive stage, three pots treated as cold stress pots were labeled and placed in the cold-water tanks in the phytotron for 10 days; the other three pots were kept at ambient temperature as a non-stress control. The temperature of air and water in the phytotron (**S4A Fig**) and ambient temperature (**S4B Fig**) was recorded five times in a day at 3-hour intervals. After 10 days of cold treatment, the cold stress pots were further placed at ambient temperature until the plants matured. A similar set of data, as well as field conditions, was collected from the labeled tillers for treated pots and six tillers from non-treated pots, and each panicle was individually harvested.

### Data analysis

Descriptive statistics-including mean, range, standard deviation, and a two-stage analytical approach were employed in R version 3.2.1 to estimate LSD (least significant difference), heritability, BLUE (best linear unbiased estimation), and pBLUP (phenotypic best linear unbiased prediction) values. Grouping of genotypes by scatter plot was performed in Minitab.

Spearman’s rank correlation coefficient was analyzed using the cor.test () function. The formula for calculating the Spearman correlation is as follows:

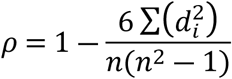

where 𝜌 = Spearman correlation coefficient; di = the difference in the ranks given to the two variables’ values for each item of the data; n = total number of observations. Principal component analysis (PCA) was performed using the prcomp() function. The grouping of germplasms was done by the hierarchical clustering method using Euclidean distance, and clustering was done by Ward’s minimum variance method (Ward.D2) via the hclust() function. All analyses were performed using R statistical software (version 4.3.1) within the RStudio integrated development environment (IDE).

## Results

### Cold tolerance at the seedling stage

Leaf discoloration (LD) and survival percentage are the two important traits for discriminating cold-tolerant lines from susceptible lines under cold stress. The mean LD score recorded at 10 days of cold stress showed a wide range of variation among the genotypes (3.7 to 8.8), and the survival rate after seven days of recovery from cold stress also varied from 0 to 37% (**Table 4**).

**Table 4:** Summary statistics of Leaf discoloration (LD) and Survival Percentage in diverse panel of rice genotypes under seedling stage cold screening.

| Parameter | LD | Survival % |
| --- | --- | --- |
| Tested entries (n = 34) | 3.7 – 8.8 | 0 – 37 |
| BRRI dhan28 (Sus. check) | 8.00 | 0.90 |
| BR1 (Sus. check) | 8.80 | 0.00 |
| BRRI dhan67 (Mod. tol. check) | 5.00 | 20.30 |
| Hbj. B. VI (Tol. check seedling) | 5.70 | 17.40 |
| p-value (Genotype effect) | $3.02 \times 10^{-8}$ | $2.04 \times 10^{-4}$ |
| CV (%) | 12.59 | 29.97 |

The susceptible checks, BRRI dhan28 and BR1, exhibited high LD values (8.0 and 8.8, respectively) coupled with very low survival rates (0.9% and 0%). In contrast, the moderately tolerant check BRRI dhan67 and the tolerant check Hbj. B. VI recorded lower LD scores (5.0 and 5.7) and higher survival percentages (20.3% and 17.4%). Analysis of variance indicated a highly significant genotype effect for both traits (p = 3.02 × 10⁻⁸ for LD; p = 2.04 × 10⁻⁴ for Survival%). The coefficient of variation was 12.59% for LD, reflecting high experimental precision, whereas Survival% showed a CV of 29.97%, indicating substantial relative variability among genotypes.

A scatter plot constructed based on combined BLUE for LD score and survival rate of 38 breeding lines, including check varieties, across the three batches presented in **Fig 1A**, and detailed data across three batches were presented in **S1 Table**. Out of 38 genotypes, two genotypes, viz., BR8907-B-1-2-CS1-4-CS2-P3-4 and BR8909-B-12-2-CS1-4-CS2-P2-3-2, obtained the lowest LD score (3.7 and 3.9) and highest survival rate (34.4-37%) in all three batches, indicating these are highly tolerant at the seedling stage. Eight genotypes (Hbj. B. VI, Bhutan, BR11318-5R-106, BR11712-4R-227, BR12266-44-11-32-5-1-1-HR10-B, BRRI dhan58, BRRI dhan89, and BRRI dhan67) formed a moderately tolerant group having LD scores of 4.9-6.5 with 17.4-31% survivability. Pictorial view of several seedling stage cold tolerant lines was shown in **Fig 1B**. Thirteen genotypes were found as moderately susceptible (LD score 5.4-7.5 with survival 6.4-15.8%), and 15 genotypes showed susceptible (LD score 5.7-8.8 with survival 0-5.7%) to cold stress at seedling stage.

**Fig 1:**
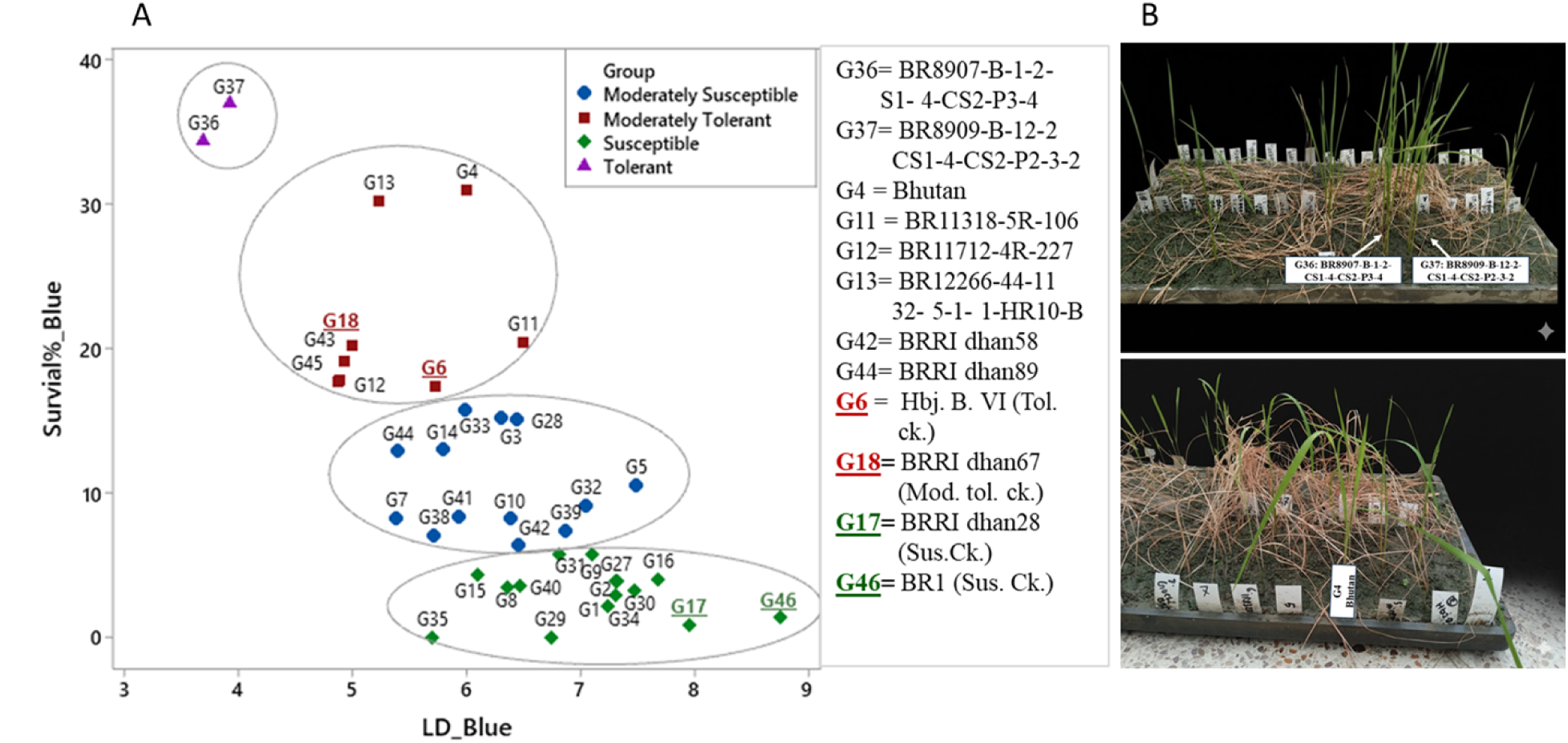
A) Clustering of genotypes based on BLUEs for LD (leaf discoloration) score and percent survival under cold water tank, B) Pictorial view of several seedling stage cold tolerant lines.

The LD score exhibited the highest variability among the evaluated traits, with genotypes fitting an approximately normal distribution (**Fig 2**). Survival percentage, in contrast, showed a narrower distribution and was skewed towards lower values. A significant and negative correlation was observed between LD score and survival percentage (Spearman’s r = -0.708, p < 0.0001), indicating that genotypes with higher LD scores showed susceptibility to cold stress and tended to exhibit lower survival percentages.

**Fig 2:**
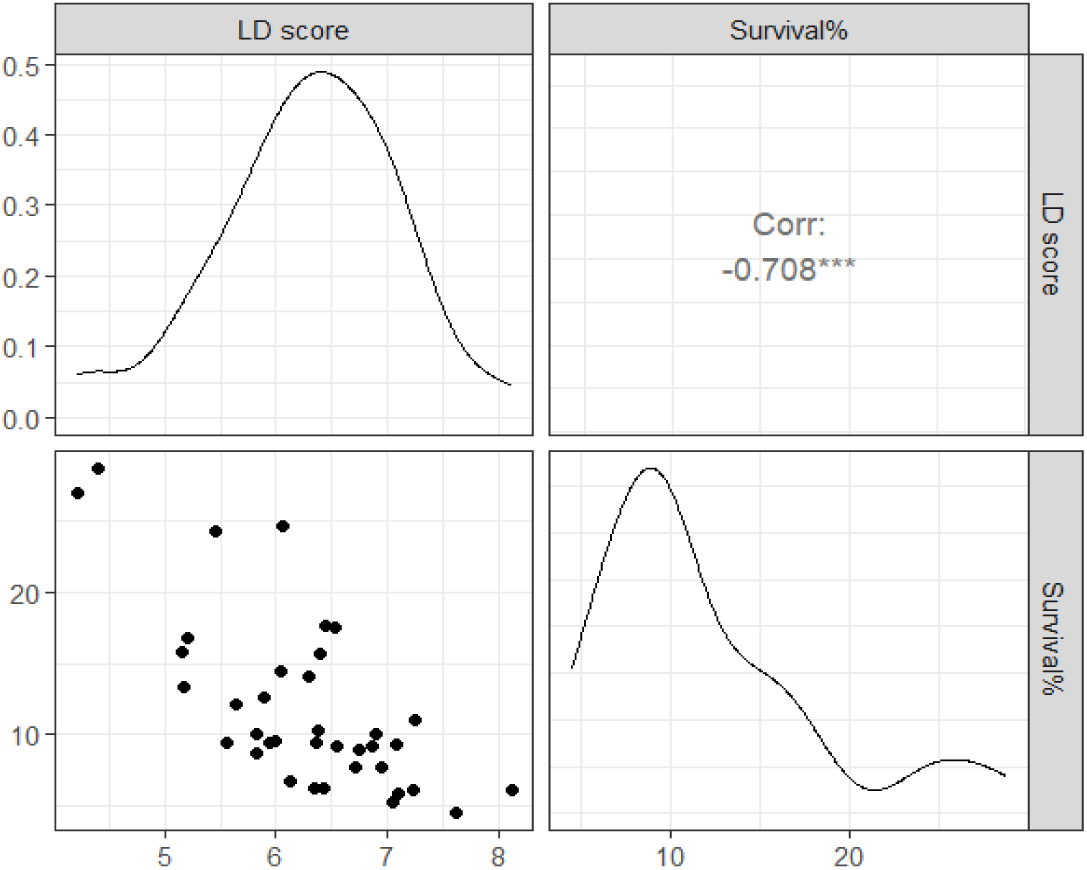
Distribution of cold tolerance phenotypes and Spearman correlation coefficients between LD score and survival% in a diverse panel of rice under cold stress in the cold-water tank.

An analysis of variance (ANOVA) was conducted to evaluate the effects of genotype, batch, and their interaction on leaf discoloration (LD) and survival percentage under cold screening conditions at the seedling stage (**Table 5**). A highly significant effect of genotype was observed both for LD and survival rate, indicating distinct LD and survival rate responses among genotypes. Batch and the genotype-by-batch interaction showed no significant interaction for LD and survival percentage, suggesting that the genotypes maintained their relative performance across different batches.

**Table 5:** Genotype-by-Batch interaction for the traits of leaf discoloration (LD) and percent survival under cold stress at the seedling stage in diverse rice genotypes.

| Trait | Source | Df | Sum Sq | Mean Sq | F value | P value |
| --- | --- | --- | --- | --- | --- | --- |
| LD | Genotype | 37 | 447.20 | 12.09 | 5.94 | <2e-16 *** |
|  | Batch | 1 | 8.50 | 8.47 | 4.16 | 0.06 |
|  | Genotype: Batch | 35 | 81.90 | 2.34 | 1.15 | 0.26 |
|  | Residuals | 380 | 773.70 | 2.04 |  |  |
| Survival % | Genotype | 37 | 32752.00 | 885.20 | 3.11 | 1.87e-08 *** |
|  | Batch | 1 | 624.00 | 624.10 | 2.20 | 0.14 |
|  | Genotype: Batch | 35 | 11954.00 | 341.60 | 1.20 | 0.21 |
|  | Residuals | 381 | 108288.00 | 284.20 |  |  |

DF: Degrees of freedom, Sum Sq: Sum of squares, Mean Sq: Mean square.

### Cold tolerance at the reproductive stage

Under cold stress conditions, all the genotypes flowered between 19 February and 28 March 2024 (**S2A Fig**), indicating that panicle initiation (PI) to booting occurred from the third week of January to the third week of February. During this period, temperatures ranged from 8.7°C to 31.4°C, with an average of 19.8°C (**S3A Fig**). Notably, between 15 and 30 January, daily averages remained below 20°C, with maximum between 18.4°C and 26.9°C, suggesting exposure to natural low temperatures during critical reproductive stages.

Genotypes flowered between 15 and 30 February, including BR12266-44-11-32-5-1-1-HR10-B, BR11303-5R-156, BR11318-5R-106, BR10317-5R-25, Bhutan, BRRI dhan74, BRRI dhan28, BRRI dhan88, IR18A1859, and TP16199, likely encountered severe cold stress at PI and booting stages. Additionally, entries flowering in early March (e.g., Rata Boro, IR83222-F11-173, Hbj. B. VI, Tepi Boro, Mineasahi, and several BR11894 derivatives) experienced moderate cold stress (19.1°C–21.3°C), particularly during booting.

In contrast, all breeding lines in non-stress plots flowered between 17 March and 10 April 2024 (**S2B Fig**). The non-stress plots recorded temperatures between 21.1°C and 25.4°C (max: 26.3°C–31.4°C) from 15 February to 10 March (**S3B Fig**), indicating favorable thermal conditions during reproductive development (PI to booting stage).

Under natural field conditions, all the phenotypic traits except vegetative stage score (VegS) showed a significant response to cold stress (**Table 4**). Although effective tiller number was significantly affected under both cold stress and non-stress conditions, it was excluded from further analysis. Specifically, in the cold stress plots, young seedlings were damaged by night birds shortly after transplanting, creating vacant spaces in the field. These gaps allowed the remaining plants to compensate by producing more tillers, resulting in a higher effective tiller number under cold stress relative to non-stress conditions. Therefore, traits like PDS, PES, SFS, plant height, days to heading, and panicle length were used for further analysis. Under phytotron conditions, these traits have a significant effect on cold stress, whereas plant height traits didn’t exhibit any significant effect on cold stress (**Table 6**). Thus, the traits that had a significant effect on cold stress were used for further analysis.

**Table 6:** Effect of cold Stress and nonstress conditions on different agronomic and cold-related traits under natural field and artificial cold screening facilities.

| Trait | Mean_cold stress | Mean_non-stress | P-value | t_statistic | Significance |
| --- | --- | --- | --- | --- | --- |
| **Natural field conditions** |  |  |  |  |  |
| Vegs | 4.92 | 4.60 | 0.14 | 1.51 | ns |
| PDS | 3.19 | 1.42 | 0.00 | 5.63 | *** |
| PES | 3.89 | 1.77 | 0.00 | 4.96 | *** |
| SFS | 5.27 | 3.54 | 0.00 | 7.29 | *** |
| Plant height (cm) | 97.07 | 104.05 | 0.00 | -3.77 | *** |
| Days to heading (days) | 163.70 | 149.71 | 0.00 | 11.85 | *** |
| Effective tiller number | 20.39 | 14.81 | 0.00 | 5.03 | *** |
| Panicle length (cm) | 21.73 | 23.93 | 0.00 | -5.97 | *** |
| **Artificial cold screening facilities** |  |  |  |  |  |
| PDS | 1.15 | 1.03 | 0.03 | 2.26 | * |
| PES | 2.35 | 1.67 | 0.00 | 4.38 | *** |
| SFS | 5.11 | 4.16 | 0.02 | 2.43 | * |
| Plant height (cm) | 97.15 | 98.44 | 0.39 | -0.87 | ns |
| Days to heading (days) | 113.52 | 108.09 | 0.00 | 7.87 | *** |
| Panicle length (cm) | 21.60 | 22.81 | 0.01 | -2.95 | ** |

\*\*\*: $P < 0.001$ , \*\*: $P < 0.01$ , \*: $P < 0.05$ , ns: Not significant ( $P\text{-value} \geq 0.05$ )
VegS: Vegetative score, PDS: Panicle degeneration score, PES: Panicle exertion score, SFS: Spikelet fertility score.

The variation in phenotypic performance of the diverse genotypes based on measured agronomic traits from non-stress to stress conditions under natural (field) and artificial cold screening facilities (phytotron) was represented by the reduction rate in **Table 7**. The lowest plant height reduction (PHR) was observed in TP16199 (G13) (-10.88% in field). BRRI dhan74 (G21) and IR100723-B-B-B-B-61 (G15) exhibited the least delay in heading date (DHD) (3.50 and 0.57 days) in field and phytotron conditions, respectively. Minimum panicle length reduction (PLR) was observed in BRRI dhan92 (G26) (-4.26% in field) and BR11712-4R-227 (G8) (-16.71% in phytotron). Whereas, the highest PHR was shown in BR10317-5R-25 (G7) (21.85% in the field). The longest DHD was observed in Mineasahi (G5) (25.50 days in the field) and BR11318-5R-106 (G10) (10.07 days in the phytotron). IR100722-B-B-B-B-11 (G14) and BR11894-R-R-R-R-329 (G17) showed the highest PLR (21.96 and 15.49%) under field and phytotron conditions, respectively.

**Table 7:** Reduction rate of the measured characters of diverse panel genotypes due to cold stress under field and phytotron conditions.

| Geno type | Designation | Field |  |  | Phytotron |  |
| --- | --- | --- | --- | --- | --- | --- |
|  |  | PHR | DHD | PLR | DHD | PLRR |
| G1 | Rata Boro | -2.00 | 12.00 | -3.17 | 10.72 | 12.42 |
| G2 | Tepi Boro | 0.35 | 18.00 | 9.27 | 4.91 | 9.12 |
| G3 | Mineasahi | -9.96 | 25.50 | -2.22 | 9.73 | 11.31 |
| G4 | Bhutan | -0.32 | 5.50 | 4.41 | 1.05 | 7.93 |
| G5 | IR83222-F11-173 (Tol. ck.) | 3.89 | 13.00 | 7.07 | 0.73 | -6.62 |
| G6 | Hbj. B. VI | 6.36 | 13.50 | -1.22 | 5.89 | -3.95 |
| G7 | IR100722-B-B-B-B-11 | 16.37 | 20.50 | 21.96 | 7.80 | 10.72 |
| G8 | IR100723-B-B-B-B-61 | 0.77 | 9.50 | 0.07 | 0.57 | 0.09 |
| G9 | BR10317-5R-25 | 21.85 | 10.00 | 15.70 | 4.69 | 14.47 |
| G10 | BR11303-5R-156 | -2.40 | 20.00 | 8.87 | 4.86 | -2.01 |
| G11 | BR11318-5R-106 | 6.43 | 11.50 | 16.00 | 10.07 | 1.24 |
| G12 | BR11712-4R-227 | 4.87 | 20.50 | 8.58 | 2.74 | -16.71 |
| G13 | BR12266-44-11-32-5-1-1-HR10-B | 3.52 | 4.50 | 19.73 | 7.65 | 16.18 |
| G14 | BR8899-14-4-1-2-2-1 | 15.39 | 12.00 | 11.97 | 5.85 | -18.35 |
| G15 | TP16199 | -10.88 | 6.00 | 12.50 | 1.69 | 6.87 |
| G16 | IR18A1859 | 7.04 | 7.50 | 21.76 | 4.91 | 2.52 |
| G17 | BRRI dhan28 (Sus. ck.) | 19.27 | 10.50 | 6.96 | 3.61 | 6.87 |
| G18 | BRRI dhan67 (Mod. tol. ck.) | 11.07 | 19.50 | 10.00 | 2.86 | 13.19 |
| G19 | BR11894-R-R-R-R-169 | 0.33 | 15.00 | 10.15 | 4.71 | 15.44 |
| G20 | BR11894-R-R-R-R-329 | 3.81 | 16.50 | 15.27 | 5.80 | 15.49 |
| G21 | BR11894-R-R-R-R-105 | 6.74 | 15.00 | 5.82 | 8.13 | 6.31 |
| G22 | BR11894-R-R-R-R-110 | 18.98 | 13.50 | 4.82 | 1.32 | 7.01 |
| G23 | BRRI dhan74 | 9.70 | 3.50 | 5.58 | 2.07 | -1.05 |
| G24 | BRRI dhan88 | 11.70 | 13.25 | 17.38 | 4.19 | 4.41 |
| G25 | BRRI dhan81 | 13.73 | 16.50 | 9.71 | 3.98 | 6.54 |
| G26 | BRRI dhan92 | 7.60 | 11.00 | -4.26 |  |  |

PHR (Plant height reduction), DHD (Delay in heading date), and PLR (Panicle length reduction)

The variation of genotypes in cold-related traits like PDS, PES, and SFS under stress and non-stress conditions in the field and phytotron was represented as paired dot plots, comparing where non-stress (blue circle) and cold stress (red triangle) scores for a given entry (**Fig 3).** A longer line indicates a greater difference between the two conditions, suggesting higher susceptibility to cold stress. Conversely, a shorter line indicates a smaller difference, implying better tolerance to cold stress. In the field, the lowest cold trait scores (indicating tolerance) were observed in Bhutan, BRRI dhan67, and BRRI dhan92 (1, PDS); BR11894-R-R-R-R-110, Mineasahi, and BRRI dhan92 (1, PES); and IR83222-F11-173 (1, SFS) (**Fig 3A**).

**Fig 3:**
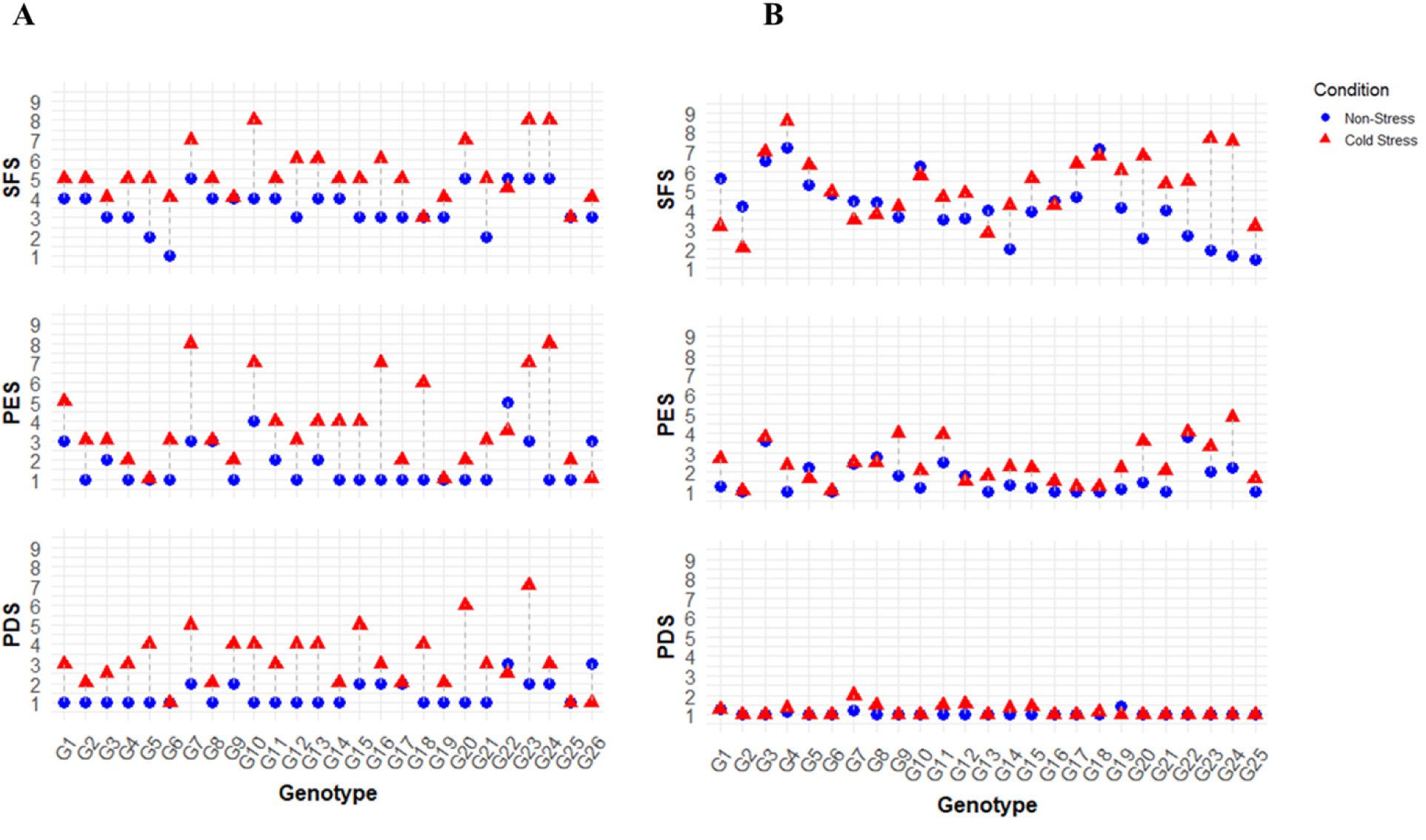
Paired dot plot showing performance of diverse rice germplasm under cold stress and nonstress conditions based on cold-related traits in A) field and B) phytotron conditions. PDS: Panicle degeneration score, PES: Panicle exsertion score, and SFS: Spikelet fertility score. Where, 1 = Highly tolerant, 3 = Tolerant, 5 = Moderately tolerant, 7 = Moderately susceptible, 9 = Susceptible.

In the phytotron, most entries exhibited low degeneration; IR83222-F11-173 and Bhutan performed the lowest PES (1), and Bhutan also showed the lowest SFS score (1; **Fig 3B)**. While BRRI dhan92 demonstrated cold tolerance in the field, this might be attributed to its longer growth duration, avoiding stress conditions. Conversely, Rata Boro and IR83222-F11-173 showed reverse results in the phytotron. SFS was higher under non-stress conditions than cold stress in the phytotron due to high-temperature stress at the reproductive stage in non-stress pots. The ambient temperature for non-stress pots at the flowering stage was 27.6-38.1℃, i.e., higher than 35℃ (**S4B Fig**). So, nonstress plots suffered from heat stress at the reproductive stage, leading to spikelet sterility.

Spearman correlation analysis was performed to elucidate the relationships among the different phenotypic variables at two different levels of cold stress (field and phytotron conditions), and the results are presented in **Table 8**. Under field, SFS exhibited a significant positive correlation with PDS and PES, indicating that spikelet fertility was associated with panicle degeneration and exsertion. A significant positive correlation was also found between plant height and panicle length. Days to heading was negatively correlated with SFS, suggesting that longer growth duration was associated with lower spikelet fertility. In the case of phytotron conditions, the traits of plant height, days to heading, and panicle length were significantly and positively correlated with each other. Days to heading showed a significant negative correlation with SFS, consistent with field observations.

**Table 8:** Spearman correlation matrix among the phenotypic variables at different levels of cold stress.

|  | PDS | PES | SFS | Plant height | Days to heading | Panicle length | Field_cold stress |
| --- | --- | --- | --- | --- | --- | --- | --- |
| PDS | 1 | 0.34 | 0.57** | -0.18 | -0.41* | -0.32 |  |
| PES | 0.08 | 1 | 0.51** | -0.14 | -0.28 | 0.08 |  |
| SFS | -0.16 | 0.18 | 1 | -0.24 | -0.21 | -0.26 |  |
| Plant height | 0.15 | -0.3 | 0.04 | 1 | 0.36 | 0.57** |  |
| Days to heading | 0.11 | -0.38 | -0.41* | 0.45* | 1 | 0.36 |  |
| Panicle length | 0.39 | -0.08 | -0.22 | 0.40* | 0.44* | 1 |  |
| Phytotron_cold stress |  |  |  |  |  |  |  |

\*\* : Significant at $P < 0.01$ , \* : Significant at $P < 0.05$
PDS: Panicle degeneration score, PES: Panicle exsertion score, SFS: Spikelet fertility score.

Principal Component Analysis (PCA) was conducted to explore the phenotypic variation and reduce the dimension among 25 rice genotypes under cold stress at the reproductive stage under the two environments (field and phytotron conditions). Under field conditions, the first two principal components (PC1 and PC2) explained 57.8% of the total variance, where PC1 accounted for 39.1% of the variability (**Fig 4A**). Whereas, under controlled phytotron conditions, PC1 and PC2 explained 31.5% and 24.6% of the total variance, summing to 56.1% (**Fig 4B**). Across both environments, G17 (BRRI dhan28), a known susceptible cultivar, consistently clustered with other susceptible genotypes on the positive side of PC1 for relevant cold traits (such as PDS, PES, SFS, PHR, and PLR in the field and PES, SFS, PLR, and DHD in the phytotron).

**Fig 4.**
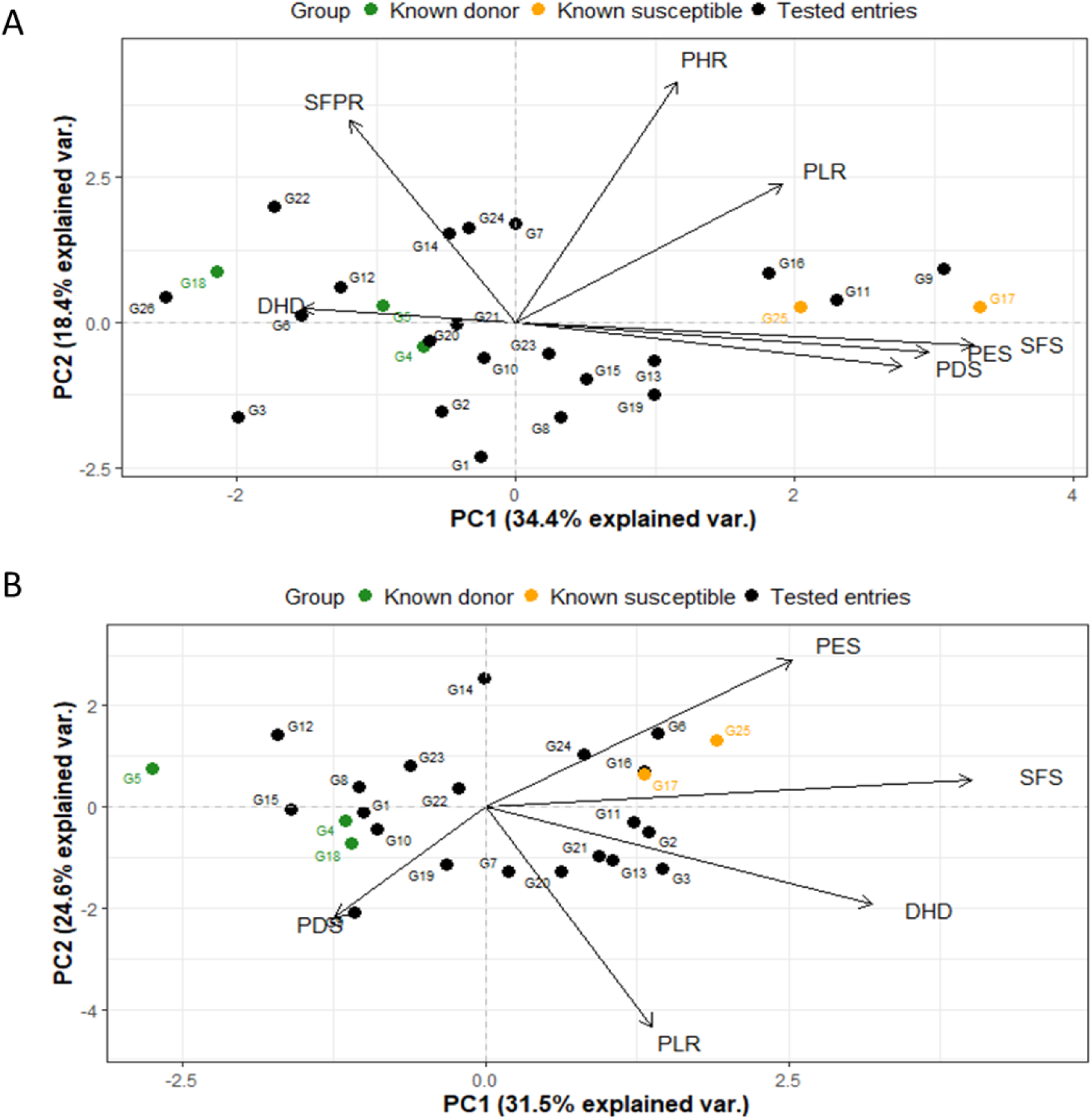
PCA analysis (PC1, PC2) showing traits and genotypes distribution regarding cold tolerance at reproductive stages in rice under A) field and B) phytotron conditions. Where, G1: Rata Boro, G2: Tepi Boro, G3: Mineasahi, G4: Bhutan, G5: IR83222-F11-173 (Tol.ck), G6: Hbj. B. VI, G7: IR100722-B-B-B-B-11, G8: IR100723-B-B-B-B-61, G9: BR10317-5R-25, G10: BR11303-5R-156, G11: BR11318-5R-106, G12: BR11712-4R-227, G13: BR12266-44-11-32-5-1-1-HR10-B, G14: BR8899-14-4-1-2-2-1, G15: TP16199,G16: IR18A1859, G17: BRRI dhan28 (Sus. Ck.), G18: BRRI dhan67 (Mod. tol.Ck.), G19: BR11894-R-R-R-R-169, G20: BR11894-R-R-R-R-329, G21: BR11894-R-R-R-R-105, G22:BR11894-R-R-R-R-110, G23: BRRI dhan74, G24: BRRI dhan88, G25: BRRI dhan81, G26: BRRI dhan92.

Under both field and phytotron conditions, PC1 is the primary descriptor of sensitivity, with sensitivity associated with positive values of SFS, PES and possibly PDS. PC2 was explained mostly by height-related parameters PHR and PLR, which may or may not be directly linked to cold stress. These traits could represent growth characteristics that are either indirectly affected by cold or simply vary independently.

However, a notable difference was observed that PDS and DHD were associated with PC1 under field conditions, but contributed more strongly to PC2 under phytotron conditions. This suggests that PDS and DHD may reflect different physiological responses depending on environmental condition. Importantly, another key discrepancy between field and phytotron conditions was DHD contribute to PC1 in the field, while PC2 under phytotron conditions. This divergence highlights the influence of environmental condition on trait expression and underscores the need to interpret PCA results within the framework of specific growing conditions.

To support and validate the PCA-based grouping of genotypes, hierarchical clustering was performed using Euclidean distance and Ward’s method under both field and phytotron conditions (S4 Fig). The resulting clusters showed strong concordance with PCA, particularly in the consistent placement of cold-tolerant genotypes in Cluster I and the susceptible check BRRI dhan28 (G23) in Cluster IV across both environments (**S5A, B Fig**).

Cluster mean values (**S2 Table**) were used to rank genotypes based on trait performance under field and phytotron conditions (**S3 Table**). The summed scores ranged from 18 to 38, with genotypes with lower summed scores across traits were classified as tolerant (18-22), while those with higher scores were considered as susceptible (33-38). The tolerant genotypes identified were Bhutan, IR83222-F11-173, BR11712-4R-227, BRRI dhan74, and Rata Boro. Genotypes which were grouped into moderately tolerant included, BR11894-R-R-R-R-110, BRRI dhan67 (Mod. tol. ck.), BR11894-R-R-R-R-105, BR11303-5R-156, TP16199, IR100723-B-B-B-B-61, BR12266-44-11-32-5-1-1-HR10-B, and BR11894-R-R-R-R-169. Susceptible genotypes were BRRI dhan88, BRRI dhan28 (Sus. ck.), IR18A1859, BR11318-5R-106, BRRI dhan81, and BR10317-5R-25.

### Performance of genotypes evaluated at both Seedling and reproductive stage

The performance of 18 common rice genotypes evaluated under both seedling and reproductive stages revealed distinct patterns of cold stress tolerance. The full spectrum of observed responses for each genotype at both developmental stages is presented in **Fig 5**. Genotypes such as Bhutan, BR11712-4R-227, BR12266-44-11-32-5-1-1-HR10-B, and BRRI dhan67 consistently exhibited moderately tolerant to tolerant phenotypes at both stages, indicating stable performance under cold stress.

**Fig 5:**
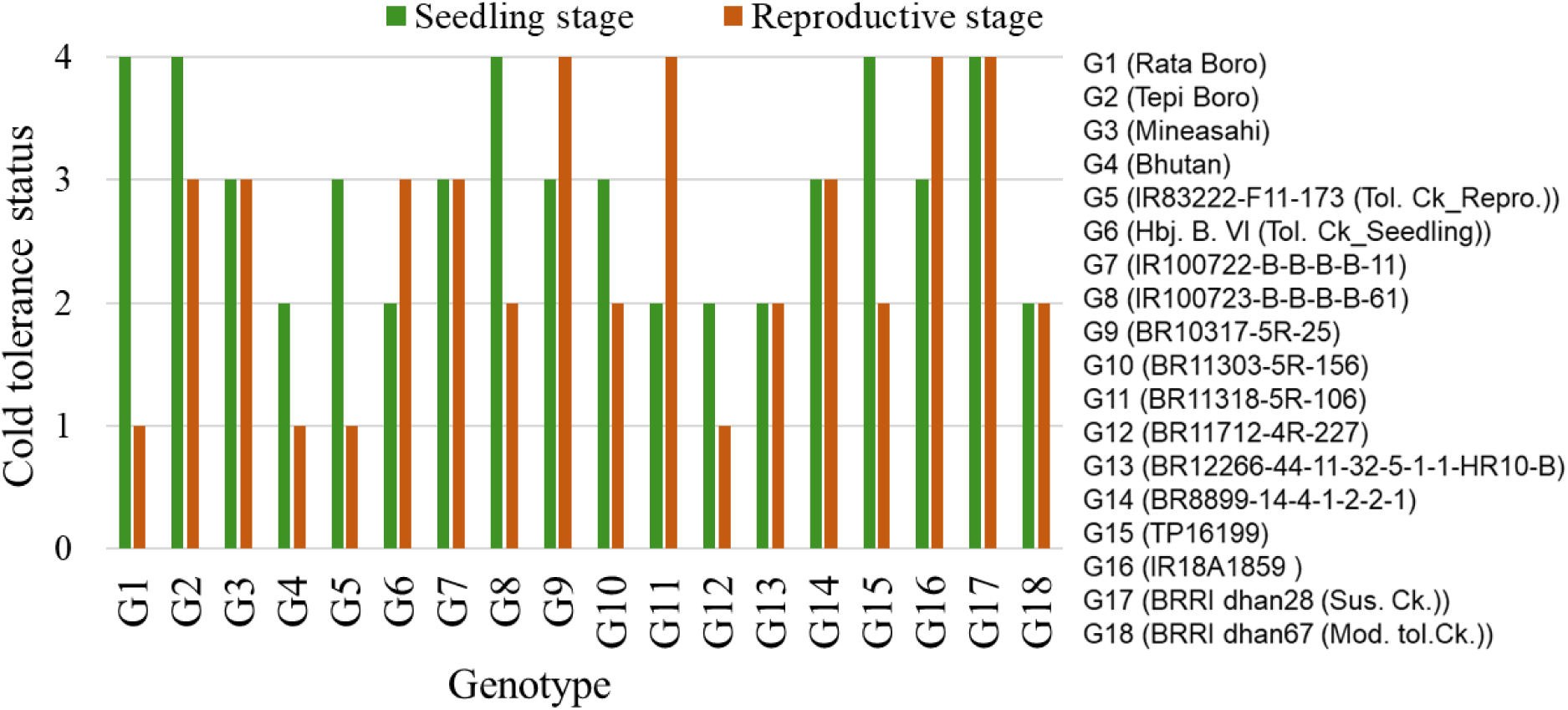
Performance of common 18 diverse rice panel under cold stress at seedling and reproductive stages. Bar heights represent the cold tolerance status for each genotype. Status was defined as: 1 = Tolerant, 2 = Moderately tolerant, 3 = Moderately susceptible, and 4 = Susceptible.

A few genotypes, including Rata Boro, BR100723-B-B-B-B-61, BR11303-5R-156, TP16199, and IR83222-F11-173, showed tolerance specifically at the reproductive stage but were moderately susceptible to susceptible at the seedling stage. In contrast, BR11318-5R-106 was moderately tolerant only at the seedling stage but susceptible at the reproductive stage. Genotypes like Mineasahi, BR100722-B-B-B-B-11, BR8899-14-4-1-2-2-1, BR10317-5R-25, IR18A1859, Tepi Boro, and the susceptible check BRRI dhan28 showed poor performance across both stages.

Spearman correlation analysis between seedling and reproductive stage results (**Table 9**) revealed a weakly positive but non-significant correlation between cold tolerance scores at the seedling and reproductive stages (r^2^ = 0.18, P = 0.479), indicating no statistically meaningful association between tolerance levels across these two developmental phases. Thus, seedling and reproductive stage cold tolerance should be considered as separate traits and quite likely under distinct genetic control.

**Table 9:** Spearman correlation for cold tolerance at seedling and reproductive stages in a diverse rice panel.

|  | Seedling Stage | Reproductive Stage |
| --- | --- | --- |
| Seedling Stage | 1 | 0.18<sup>NS</sup> |
| Reproductive Stage | 0.18<sup>NS</sup> | 1 |

Cold tolerance was coded as: 1= Tolerant, 2= Moderately tolerant, 3= Moderately susceptible, 4= Susceptible.

A PCA was employed to investigate the relationship among cold stress-related traits at both the seedling (LD, survival percentage) and reproductive stages (PDS, PES, SFS, PHR, DHD, PLR) across 18 common rice genotypes, under field and phytotron conditions (**Fig 6**).

**Fig 6:**
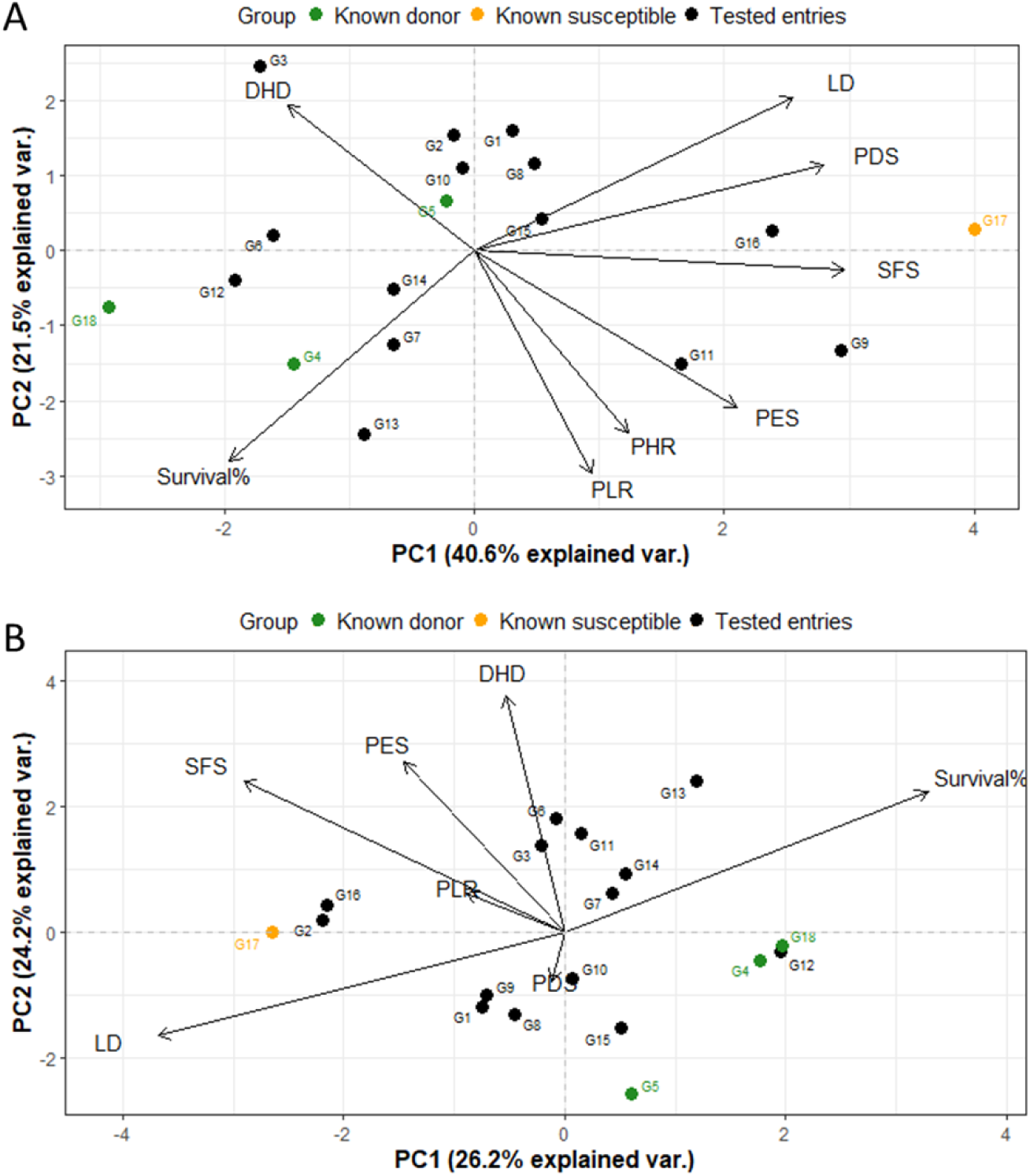
PCA biplot of 18 rice genotypes based on cold stress-related traits at seedling and reproductive stages under A) field and B) phytotron conditions. Where, G1: Rata Boro, G2: Tepi Boro, G3: Mineasahi, G4: Bhutan, G5: IR83222-F11-173 (Tol.ck), G6: Hbj. B. VI, G7: IR100722-B-B-B-B-11, G8: IR100723-B-B-B-B-61, G9: BR10317-5R-25, G10: BR11303-5R-156, G11: BR11318-5R-106, G12: BR11712-4R-227, G13: BR12266-44-11-32-5-1-1-HR10-B, G14: BR8899-14-4-1-2-2-1, G15: TP16199, G16: IR18A1859, G17: BRRI dhan28 (Sus. Ck.), G18: BRRI dhan67 (Mod. tol.Ck.).

Under field conditions (**Fig. 6A**), the first two principal components (PCs) accounted for 62.1% of the total variation, with PC1 contributing 40.6%. PC1 had a greater contribution of LD, PDS, and SFS, higher values of these traits corresponding to cold susceptibility. On the other hand, PLR, PHR, DHD and survival percentage contributed to PC2. Thus, a combination of seedling and reproductive traits contributed to each of the first two principal components.

Under phytotron conditions (**Fig. 6B**), PC1 and PC2 explained 50.4% of the total variation (PC1: 26.2%, PC2: 24.2%). PC1 was mostly dominated by seedling traits LD and survival%, with some contribution from SFS, while PC2 was mostly controlled by reproductive traits DHD, PES and SFS. Thus, phytotron data shows distinct partitioning of PC1 and PC2 between seedling and reproductive-stage traits, supporting the hypothesis that seedling and reproductive-stage tolerance are separate traits having different genetic control.

## Discussion

Cold stress adversely affects rice productivity, with its varying effects depending on growth stage as well as the severity and duration of exposure. Rice plants are susceptible to cold damage at any developmental phase - including germination, seedling, vegetative, reproductive, and maturity stages [10,19]. This study investigates a diverse set of rice genotypes under cold conditions during the seedling and reproductive stages, aiming to elucidate stage-specific vulnerabilities and to reinforce the importance of comprehensive cold screening at all growth stages to unravel the complex resilience pathways to cold stress.

Cold damage at the seedling stage manifests primarily as leaf yellowing due to inhibited chlorophyll synthesis, stunted growth, and eventual seedling mortality. These physiological disruptions often reduce photosynthetic efficiency and elevate respiration, leading to assimilate shortages that impair tissue development and halt growth. LD is widely used to differentiate cold-tolerant genotypes from susceptible ones; however, it alone does not fully capture the complexity of seedling-stage cold tolerance [17].

Therefore, survival rate is also considered a critical metric for identifying promising donor genotypes [19]. In this study, 38 diverse rice genotypes were evaluated for seedling stage cold tolerance across three batches (B1, B2, B3) using both LD and survival percentage. Two genotypes, BR8907-B-1-2-CS1-4-CS2-P3-4 (G36) and BR8909-B-12-2-CS1-4-CS2-P2-3-2 (G37) consistently demonstrated superior tolerance across all batches, with the lowest LD scores (3.7 and 3.9) and highest survival rates (34.4-37%). These lines, derived from three-way crosses involving donor parents 89010-TR1232-4-1 and CUNJING15 under BRRI dhan29 and BRRI dhan28 backgrounds, respectively, represent valuable resources for breeding programs. The performance of the tolerant and susceptible varieties aligned with their expected tolerance categories (Fig 3), justifying the tolerance levels of G36 and G37. Also six genotypes-BR11318-5R-106 (G11), BR11712-4R-227 (G12), BR12266-44-11-32-5-1-1-HR10-B (G13), Bhutan (G4), BRRI dhan58 (G43), and BRRI dhan89 (G45) were classified as moderately tolerant, showing LD scores of 4.9-6.6 and survival rates of 14.1-31%. Among these, Bhutan and BR12266-44-11-32-5-1-1-HR10-B showed relatively high survival (24.3-24.7) despite moderate LD, indicating possible recovery or stress avoidance mechanisms. This partial independence of LD and survival percentage suggests distinct physiological pathways governing cold tolerance for these two traits in some genotypes. Suh et al. (2013) reported LD scores ranging from 3 to 8 (mean value of 5) among 23 elite cultivars grown under cold-water irrigation, reinforce LD as a visual assessment of damage, while survival percentage offers direct measure of resilience. The strong negative correlation observed between LD score and survival percentage (Spearman’s r = -0.708, p < 0.0001) indicates these two parameters are closely related and can, at least partially, substitute for each other in assessing seedling-stage cold tolerance. Genotypes with low LD and high survival are reliably tolerant, while outliers may possess unique genetic and physiological mechanisms worth further investigation.

Reproductive-stage cold stress in rice was evaluated under field and phytotron conditions to assess genotype performance across environments. Screening at young microspore stage using cold air, water and natural winter conditions enabled consistent evaluation [22]. The trait values were normalized as reduction rates (e.g., PHR, PLR, DHD) relative to controls [15,19], along with visual injury scores (PDS, PES, and SFS), to quantify sterility and panicle development failure (**Table 7**; **Fig 3A, B**). These remain standard indicators for reproductive stage cold tolerance [19,23].

Significant phenotypic variation among the genotypes across environments revealed diverse cold response mechanisms. Genotypes like Bhutan, IR83222-F11-173, Rata Boro, and BRRI dhan74 showed consistently low reduction rates and injury scores, suggesting physiological resilience to cold stress. By contrast, the genotypes TP16199, BR11303-5R-156, and BR12266-44-11-32-5-1-1-HR10-B showed moderate cold tolerance scoring and moderate level of reduction in the studied traits, while several genotypes (BR11894-R-R-R-R-110, BR11303-5R-156, Rata Boro, IR83222-F11-173, and BRRI dhan67) exhibited delayed heading (12-20.50 days) but maintained low PHR, PLR, PDS, PES, and SFS, suggesting escape of cold stress in these genotypes due to the delayed heading. However, the opposite result was reported by Suh et al. (2010) [7,24]. Thus, DHD alone may not be a reliable measure for screening reproductive-stage cold-tolerant rice [25].

Correlation analysis revealed that growth reduction traits (PHR, PLR, DHD) were positively associated, as were traits related to injury scores (PDS, PES, SFS) (**Table 8**), while negative correlations were observed between injury score and growth reduction traits, indicating that greater reduction in growth traits aligns with higher injury. This observation is consistent with the findings [19,24] that cold stress reduces culm length and panicle development, ultimately impairing spikelet fertility. PCA distinguished the genotype responses, with DHD contributing to PC1 in field condition implying delayed flowering as a cold escape strategy. Under phytotron condition, DHD explained by PC2. This distinct shift in DHD’s contribution between the two environments highlights a significant Genotype×Environment interaction, suggesting that the phenotypic traits governing cold stress variability differ based on growth conditions (**Fig 4**).

Comparable studies [15] have validated the Average Tolerance Index (ATI) for rice germplasm assessment under cold stress at early growth stages emphasizing genotype × environment interactions in trait expression.

Hierarchical clustering and trait-based ranking demonstrated stable performance of genotypes such as IR83222-F11-173, Bhutan, Rata Boro, and BRRI dhan74 across both environments, suggesting robust cold tolerance (**S5 Fig, S3 Table**). This validates previous claims that cultivars tolerant under controlled cold conditions also perform well under field stress [23], supporting phytotron screening as a reliable proxy. Eighteen genotypes evaluated at both seedling and reproductive stages showed a weak and non-significant correlation between the two stages indicating different genetic mechanisms governing cold tolerance at each stage. This is supported by studies showing distinct QTLs and physiological responses control tolerance at seedling (e.g., *qCTS12.1*) and reproductive stages (e.g., *qPSR10*) [5][9]. For example, early-stage traits such as germination percentage, shoot and root length, and coleoptile length are strongly intercorrelated, but non-significant and negligible positive association between the germination and early seedling establishment [15]. Together these results emphasize the need to screen germplasm for cold tolerance at both stages, rather than assuming tolerance at one stage will predict tolerance for another.

PCA combining both developmental stages under field conditions indicated seedling and reproductive traits may share similar physiological pathways, supporting the hypothesis that certain genotypes possess integrated tolerance mechanisms operating across the developmental stages. This is consistent with recent studies suggesting that cold stress responses involve shared signaling pathways - such as ABA-mediated regulation, ROS detoxification, and membrane stabilization - active during both vegetative and reproductive phases [26]. However, under phytotron conditions, seedling and reproductive traits contributed to different components, supporting partial independence and stage-specific genetic control. These findings confirm that while cold tolerance may involve shared pathways, independent regulation is also evident, necessitating stage-specific screening for accurate genotype classification.

## Conclusions

This study successfully identified rice genotypes with differential responses to cold stress at both seedling and reproductive stages through multi-environment and multi-stage screening. The use of artificial cold-water tanks for seedling-stage evaluation and combined natural and controlled conditions for reproductive-stage assessment enabled precise phenotyping across developmental phases. Genotypes such as BR8907-B-1-2-CS1-4-CS2-P3-4 and BR8909-B-12-2-CS1-4-CS2-P2-3-2 consistently exhibited seedling-stage cold tolerance, while Rata Boro, BR100723-B-B-B-B-61, BR11303-5R-156, TP16199, and IR83222-F11-173 demonstrated reproductive-stage cold tolerance. Notably, Bhutan, BR11712-4R-227, and BR12266-44-11-32-5-1-1-HR10-B showed moderate to high tolerance across both stages, indicating potential for broad-spectrum cold tolerance. Conversely, several genotypes including BRRI dhan28, BR10317-5R-25, and IR18A1859 were found to be susceptible at both stages.

Principal Component Analysis revealed that under field conditions, seedling and reproductive traits contributed jointly to variation in both PC1 and PC2, suggesting physiological integration. However, under phytotron conditions, PC1 mostly captured by seedling-stage traits while PC2 was exclusively defined by reproductive-stage traits, indicating unique genetic regulation when environmental variability is minimized.

## Supporting information

**S1 Fig.** Temperature profile of batch 1, 2 and 3 in cold water tank.

**S2 Fig**. Dot plot showing flowering window of the diverse panel under A) early sown cold stress plot (CS) and B) regular Boro season (NS) plot in field.

**S3 Fig.** Maximum and minimum temperature of field conditions under A) cold stress, B) Non stress conditions.

**S4 Fig**. Temperature profile of air and water in artificial cold screening facilities (CSF) of A) cold stress, and B) ambient air temperature.

**S5 Fig**. Dendrogram showing the hierarchical clustering of rice genotypes based on phenotypic traits under (A) field and (B) phytotron conditions with cold stress at the reproductive stage.

**S1 Table.** Phenotypic performance in term of LD score and survival rate (%) of 38 rice genotypes after cold stress (13°C) at seedling stage

**S2 Table.** Cluster mean of different cold traits in the clusters formed at different cold levels using Euclidian distance matrix

**S3 Table**. Ranking of diverse rice germplasm for their relative cold related traits at reproductive stage in a cluster analysis using Euclidian distance matrix

## Acknowledgments

The authors wish to thank Md. Abu Syed, Ferdous Prince, Md. Rafiqul Islam, Sharmistha Ghosal, Ripon Roy, Ratna Rani Majumdar, Md. Abu Shama, Fahima Khatun for technical assistance and support in various aspects of this work.

## Author Contributions

Conceptualization: PSB JDP AKMAI FA. Data curation: FA PSB. Formal analysis: FA. Funding acquisition: PSB KMI, Investigation: PSB JDP AKMAI. Methodology: PSB JDP AKMAI FA Project administration: KMI PSB. Supervision: PSB JDP AKMAI. Drafting: FA. Review & editing: JDP PSB AKMAI MSR MRI.

